# SANTON: Sequencing Analysis Toolkits for Off-target Nomination

**DOI:** 10.1101/2025.05.09.653082

**Authors:** Huan Qiu, Ruijia Wang, Shu Wang, Aymen Maqsood Mulbagal, Eric Anderson, Huanying Gary Ge

## Abstract

**Background:** Genome-wide off-target nomination and screening sequencing methods, such as synthetic oligonucleotide-based sequencing and whole genome-based sequencing, evaluate the precision and safety of gene-editing technologies. However, there remains a lack of comprehensive bioinformatics tools for analyzing the sequencing data generated by off-target nomination assays for various gene editors, including Cas9, Cas12a, and base editors.

**Results:** We introduce a Sequencing Analysis Toolkits for Off-target Nomination (SANTON) for the identification and quantification of potential off-target sites using datasets generated from synthetic oligonucleotide-based sequencing and whole genome-based sequencing (e.g., Digenome-seq) methods. By applying SANTON to a Cas9 treated oligo synthesization-based sequencing data, a comprehensive set of potential off-target sites are evaluated in which the top off-target sites displayed highly constituency with a published study using *in vivo* method (e.g, GUIDE-seq). Utilizing SANTON to previously published Digenome-seq datasets, we identified more potential off-target regions than previous studies, including ones showing significantly higher cleavage level than the on-target site. To the best of our knowledge, SANTON is the first public package capable of analyzing synthetic oligonucleotide-based sequencing data and the first to specific optimized for the analysis of whole genome-based sequencing analysis for Cas12a and base editors. We demonstrated the capability of SANTON to effectively analyze complex cleavage patterns by a diverse editing system.

**Conclusions:** SANTON provides a powerful computational solution to support genome editing research, facilitating more reliable and comprehensive off-target profiling.

## Background

The CRISPR-Cas9 technology has revolutionized the field of genome editing, providing a versatile and efficient tool for introducing precise modifications to the genomes of various organisms [1–3]. Despite its widespread use, a significant challenge remains in the identification and quantification of off-target effects that can lead to unintended genetic alterations with potentially deleterious consequences. These challenges limit the safe and effective application of CRISPR-based technologies in both research and therapeutic contexts.

Various efforts have been made in the development of next-generation sequencing (NGS) assays to address this issue. Multiple genome-wide off-target identification and screening sequencing techniques have been developed. For example, Digenome-seq is a prominent method for *in vitro* off-target (OT) detection that leverages whole-genome sequencing to identify potential off-target sites [4]. However, the current available tool does not fully support analysis of data treated by Cas12a and adenine base editors (ABEs) and cytosine base editors (CBEs), which limits its applicability in diverse genome editing studies. Synthetic oligonucleotide-based sequencing [5] is another powerful *in vitro* approach that can provide comprehensive off-target profiles by integrating biochemical assays with next-generation sequencing. Despite its potential, there is no publicly available tool for analyzing any oligo synthesization and enrichment based sequencing data, creating a significant gap in the bioinformatics resources available for off-target profiling. The lack of comprehensive tool available for analyzing all different types of sequencing data from both Cas9, Cas12a, and base editing systems, limit the broader application and optimization of genome editing technologies.

Here, we present SANTON, a novel bioinformatics package designed to address this critical requirement. SANTON offers a unified platform for the identification and quantification of potential off-target sites resulting from diverse gene editing systems, including Cas9-mediated, Cas12a-mediated, and base editing platforms. Applying SANTON to available sequencing datasets shows a high level of consistency with prior studies and reveals more potential off-target regions, demonstrating its reliability and power in genome-editing assessments.

### Implementation

SANTON consists of two major modules (Fig. 1): the synthetic oligonucleotide-based sequencing module (SOS module), the equivalent of which has not been publicly available to our knowledge, and the whole genome-base sequencing module (WGS module), which introduces notable enhancements over the existing implementation [6].

**Fig 1.**
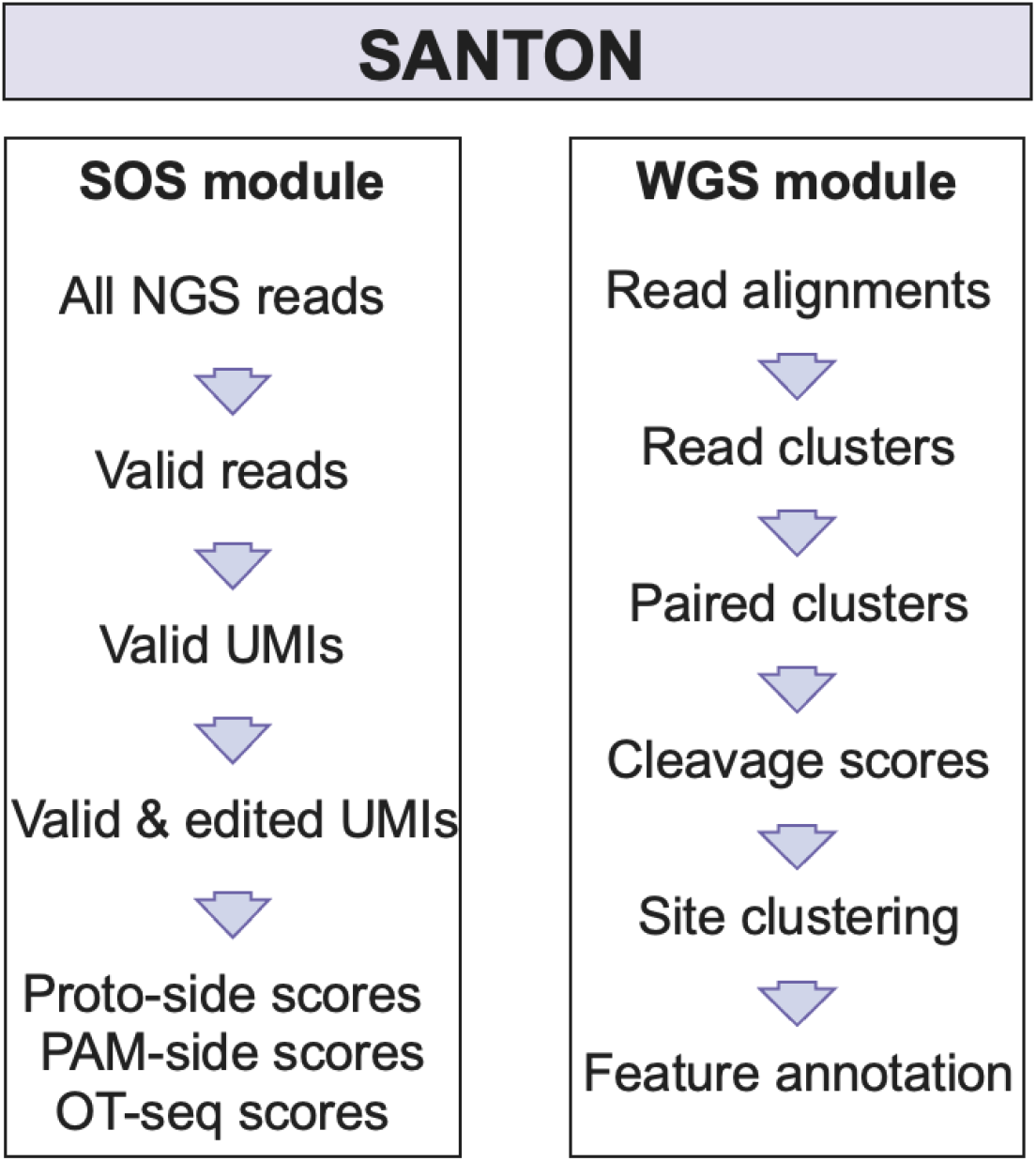
Overview of SANTON package and high-level summary of workflow under SOS and WGS modules.

### Workflow of data analysis in SOS module

#### Structure of sequencing library and sequencing reads

Each predicted off-target together with its flanking regions are embedded in a structure depicted in Fig. 2. Constant regions 1, 2, and 3 are located at 5’ upstream to the proto-side of off-target sequence, while constant regions 4, 5, and 6 are located at 3’ downstream of its pam-side. Unique Molecular Identifier (UMI) sequences, of which length is 11 base pairs (bps), are randomly inserted between constant region 1 and constant region 2, as well as between constant region 5 and constant region 6. A 14 bps Barcode sequences which is unique for each predicted off-target sites are inserted between constant region 2 and constant region 3, as well as between constant region 4 and constant region 5. *In vitro* cleavage inside of the off-target leads to proto-side fragment and pam-side fragment. After adaptor ligation to the cleavage end, these two types of fragments are subjected to NGS sequencing.

**Fig 2.**
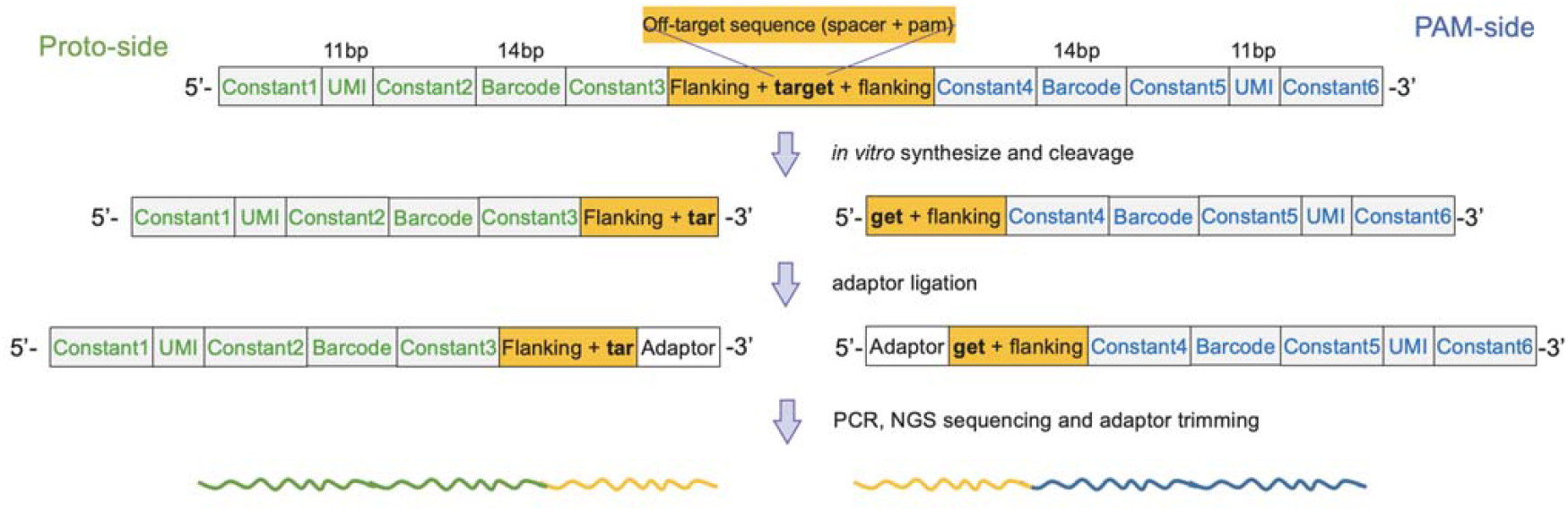
Schematic illustration of template design for synthetic oligonucleotide-based sequencing and consequence by *in vitro* cleavage. The building blocks of library sequence for each off-target candidate is depicted in order from 5’-end to 3’-end. UMIs are 11-bp oligo nucleotides located between constant region-1 and -2 in proto-side and between constant-5 and -6 regions. Barcodes are 14-bp oligo nucleotides located between constant region-2 and -3 in proto-side and between constant-4 and -5 regions. Following *in vitro* cleavage, library templates corresponding to true off-target are split into two fragments with breakpoint located within off-target region (yellow). The breakpoints are ligated with sequencing adaptor for NGS sequencing.

#### Quality Control and Filtering of Sequencing Reads

The raw NGS reads were initially filtered and consolidated based on UMIs (Fig. S1, Additional file 1). Proto-side reads containing intact constant regions-1, -2, and -3 or pam-side reads with intact constant regions-4 and -5 were retained (Fig. S1, Additional file 1). The barcode sequence of each read was extracted, and only those matching the pre-designed list were kept as valid reads (Fig. S1, Additional file 1). To minimize PCR biases, reads with identical UMI and barcode combinations from the same side were consolidated (Fig. S1, Additional file 1). The validated UMI were then used for downstream editing and quantification analysis.

#### Editing Quantification on Candidate Off-Target Sites

The fragment of putative off-target regions was extracted as sequence downstream of constant-3 in proto-side UMIs and upstream sequence to constant-4 in pam-side UMIs (Fig. 2). The extracted fragment was aligned against its corresponding reference sequences using the Smith-Waterman algorithm implemented in pairwise2 function from BioPython [7].

For Cas9-mediated cleavage, proto-or pam-side UMIs were considered edited if the edge of alignment is within ±3bp window centering the expected cut site position, and the total number of mismatches plus gaps are less than 3 (Fig. 3A). Similar criteria are applied to other editing systems with tweaked cleavage windows as well as cutoffs for mismatches and gaps (Fig. 3A). Additional requirements are applied to Base editors: 1) For ABE, adenine-to-guanine conversion within the 5’-upstream region of nick site on the proto-side was required (Fig. 3B). 2) For CBE, because cytosine-to-thymine conversions at PAM side are removed by Uracil-specific excision reagent (USER) during library preparation when double-stranded break is created, there are no nucleotide conversions in sequenced UMIs. For this reason, we require a cytosine nucleotide located at the end of pam-side read alignment or its immediate 5’-upstream position (Fig. 3C). For each off-target template, the edited fragments were quantified by counting the associated unique UMIs.

**Fig 3.**
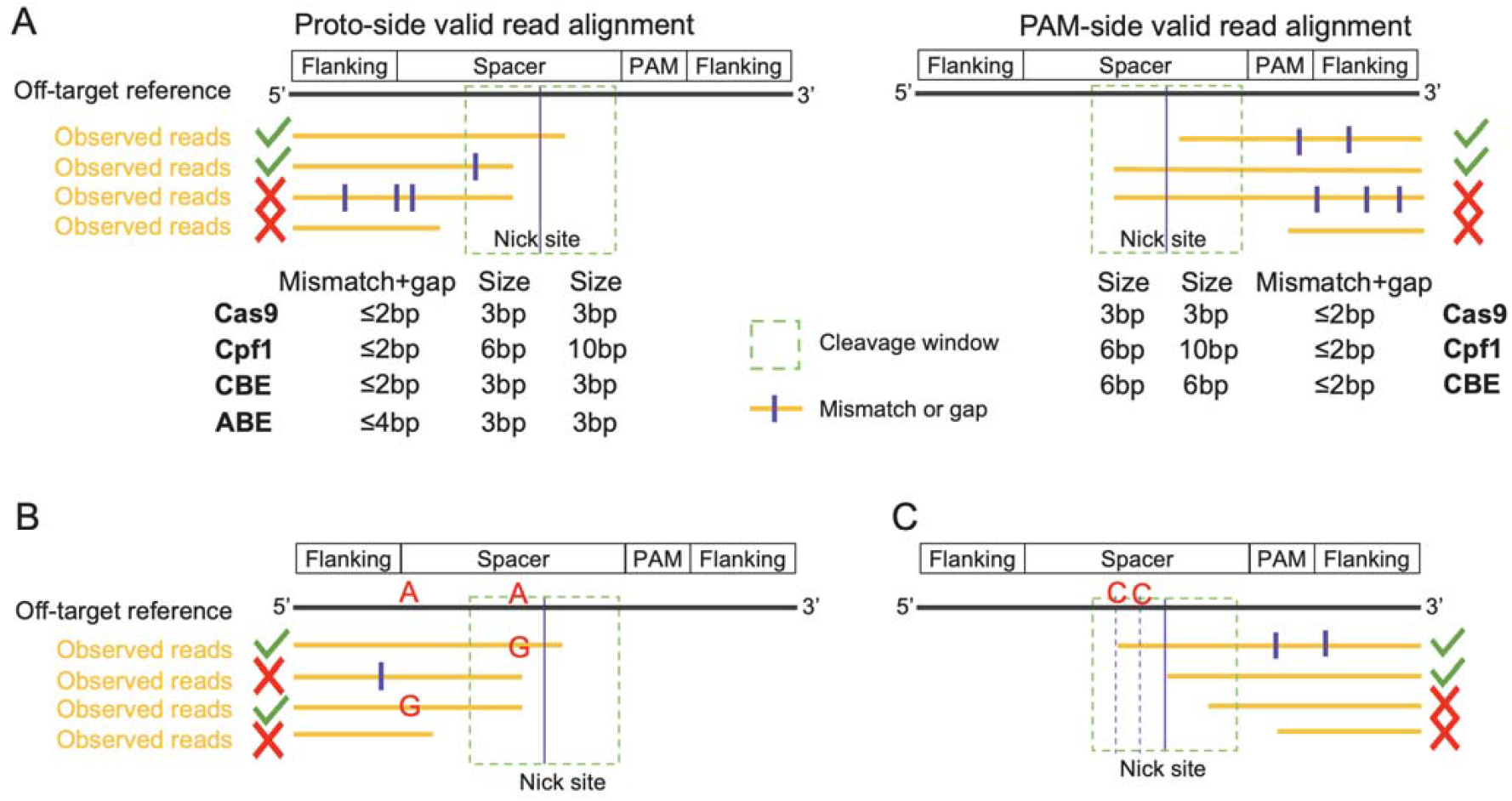
Definition of edited synthetic oligonucleotide-based sequencing UMI. (A) Allowed cleavage window size and mismatches/gaps in alignments for each editing system. Black and yellow horizontal lines stand for off-target reference sequence and observed UMIs, respectively. Vertical tick stands for mismatch or gap. Green checkmarks stand for valid UMIs that meet all the requirements of edited UMI definition. Red crosses stand for valid UMIs that fail to meet those requirements. (B) Additional requirement for edited UMI in ABE-treated data. Nucleotide conversion (A>G) must present in the region 5’-upstream to nick site on the proto-side. (C) Additional requirement for edited UMI in CBE-treated data. A cytosine nucleotide must present at the end of pam-side alignment or the 5’-upstream position next to it.

#### Off-Target (OT) scoring

To quantify the potential editing level on each candidate off-target template, OT scores are calculated for proto- and pam-side separately, in which the edited UMI count for off-target candidates is divided by the on-target region within the same panel. The final OT score was averaged from proto-side and pam-side scores for all data types except ABE-treated data. For the latter, the proto-side score was treated as OT score [5], as the editing outcome (A>G conversion) is only observable in the edited proto-side UMIs.

### Workflow of data analysis in WGS module

#### Identification and quantification of cleavage sites in single-gamma scenario

To identify the potential cleavage sites, the 5’ alignment position are first extracted from the genomic mapped reads from input BAM file using pysam (https://pysam.readthedocs.io/en/latest/api.html) (Fig. 4A). The associated forwardly-or reversely-mapped reads are retained as supporting reads and are grouped into clusters separately based on their shared 5’-end alignment positions (Fig. 4B). Previous studies [4, 6] used gamma (G) to define different cleave patterns, with 0 value stands for blunt-end, greater than zero values for overlapping, and less than zero values for gapped alignments (Fig. 4C). Paired forward- and reserve-read clusters that correspond to different gamma values are constructed based on relative distance between the 5’-end position of their read alignment using BEDTools [8]. The cleavage score is calculated following equation formulated in previous study [4, 6] to evaluate the sum of weighted supporting ratio for each pair of read clusters (Fig. S2, Additional file 1).

**Fig 4.**
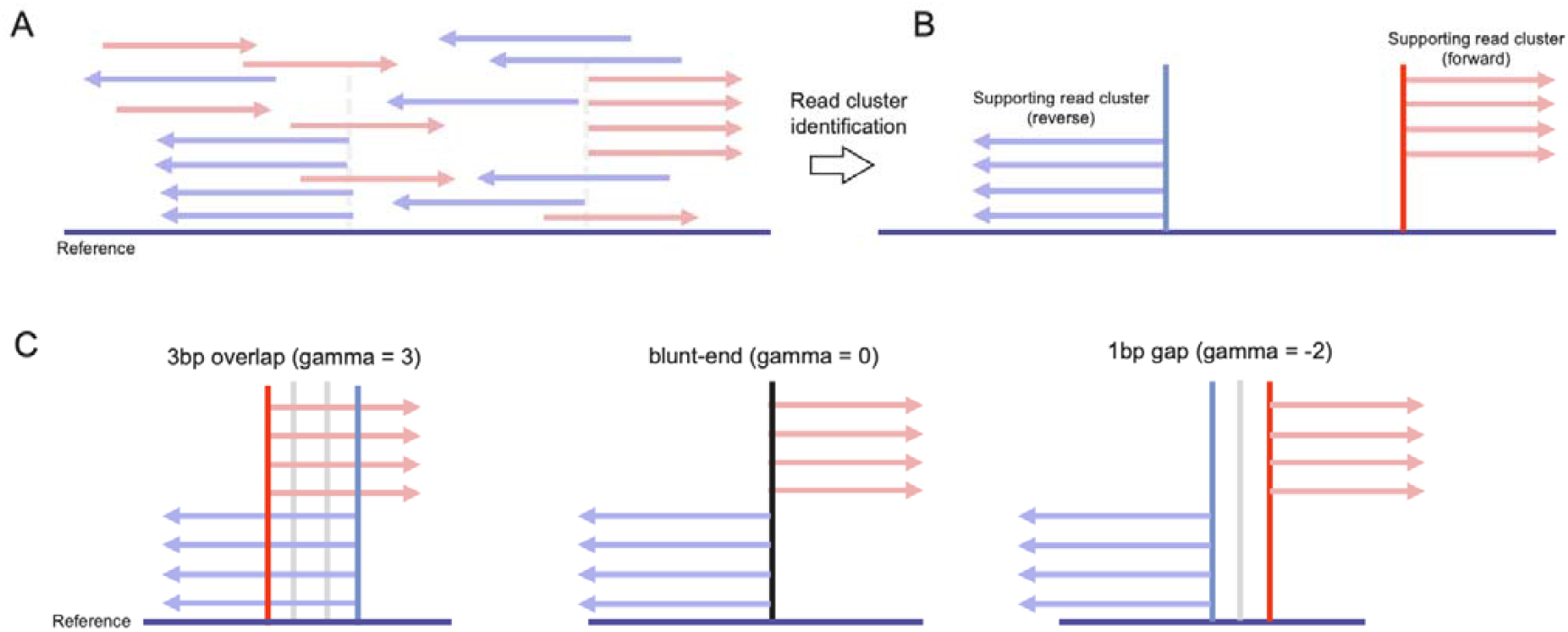
Schematics of typical whole genome-based sequencing (e.g., Digenome-seq) read mapping patterns. (A) Simplified illustration of reads mapped to reference genome sequence. Horizontal bars stand for reference genome sequence. Arrows in salmon and blue colors stand for forwardly- and reversely-mapped reads, respectively. (B) Simplified illustration of supporting read clusters that contain multiple reads with shared 5’ alignment positions. (C) Example of different reads mapping patterns formed by read cluster pairs. Horizontal bar and arrows of different colors stand for genome reference and mapped reads as mentioned earlier.

#### Identification and quantification of cleavage sites in multi-gamma scenario

The distance between nick site and editable nucleotide (cytosine for CBE or adenine for ABE) varies across off-target region and often differs from on-target region. To identify DNA cleavage sites caused by base editors (BE), paired forward-reverse read clusters with overlaps ranging from one to 19 base pairs were considered (gamma = 1∼19), provided there is one or more editable nucleotide within the 5-bp flanking region on the distal side of off-target region. For Cas12a cleavage, paired forward-reverse read clusters with gamma ranging from -10 bp to +10 bp were considered. In case of base editor and Cas12a, more than one cleavage sites are typically aggregated inside or around an off-target region (see Fig. S3, Additional file 1 for examples). To remove redundancy, clusters are inferred by grouping cleavage sites that are ≤ 5bp apart using single-linkage clustering method. The cleavage site with the highest score is selected as representative of its cluster (Fig. S4, Additional file 1).

## Results

### Nomination of off-target cleavage sites of EMX-1 gRNA using internal data with SOS module

To demonstrate the use of SANTON package on the synthetic oligonucleotide-based sequencing data, we designed a custom panel targeting the *in-silico* predicted off-target regions of EMX1 gRNA mediated by Cas9 encompassing 81,855 oligo templates. The resulting library was built and sequenced using the Illumina Mi-Seq platform leading to 13.6 million reads. Analysis of this data using SOS module (Additional file 3) shows that all templates are covered by sequencing with 81,613 (>99%) having ten or more UMIs (Fig. S5, Additional file 1). The on-target region displays high sequence coverage (>1,000 UMIs) and exceedingly high edited ratios (>90%) (Fig. S5, Additional file 1). By leveraging high-resolution UMI-based sequencing data, SANTON enables more precise quantification of off-target events, capturing potential off-target sites that might be overlooked by other approaches. A total of 521 off-target regions shows a level of cleavage close or higher the on-target (score >0.1), which includes most (86.7%) of the off-target regions that reported from previous *in vivo* EMX1 off-target study using GUIDE-seq method [9] (Table S1, Additional file 2). The high consistency between SANTON’s and in vivo sequencing method on the off-target sites display its reliability and accuracy. Of the seven off-target regions with score >1, two are identified by this GUIDE-seq experiment [9]. These off-target regions are well supported as exemplified in the off-target region (chr8:127788995-127789018) with score=1.09 (Fig. 5A). A total of 6,135 out of 6,352 (96.58%) UMIs representing 144 alignments mapped to the proto-side of this region are edited UMIs that support cleavage within the defined window. The five most abundant alignments account for the majority of the UMIs and are shown in Fig. 5A. A similar pattern can be also observed on the PAM-side of the same region, in which the most abundant alignment species from the two sides often complement each other, collaborating cleavages at the corresponding nucleotide positions (Fig. S6, Additional file 1). The above results confirm the robustness of SANTON in off-target analysis, providing a systematic and scalable framework that improves the precision and reproducibility of synthetic oligonucleotide-based sequencing studies.

**Fig 5.**
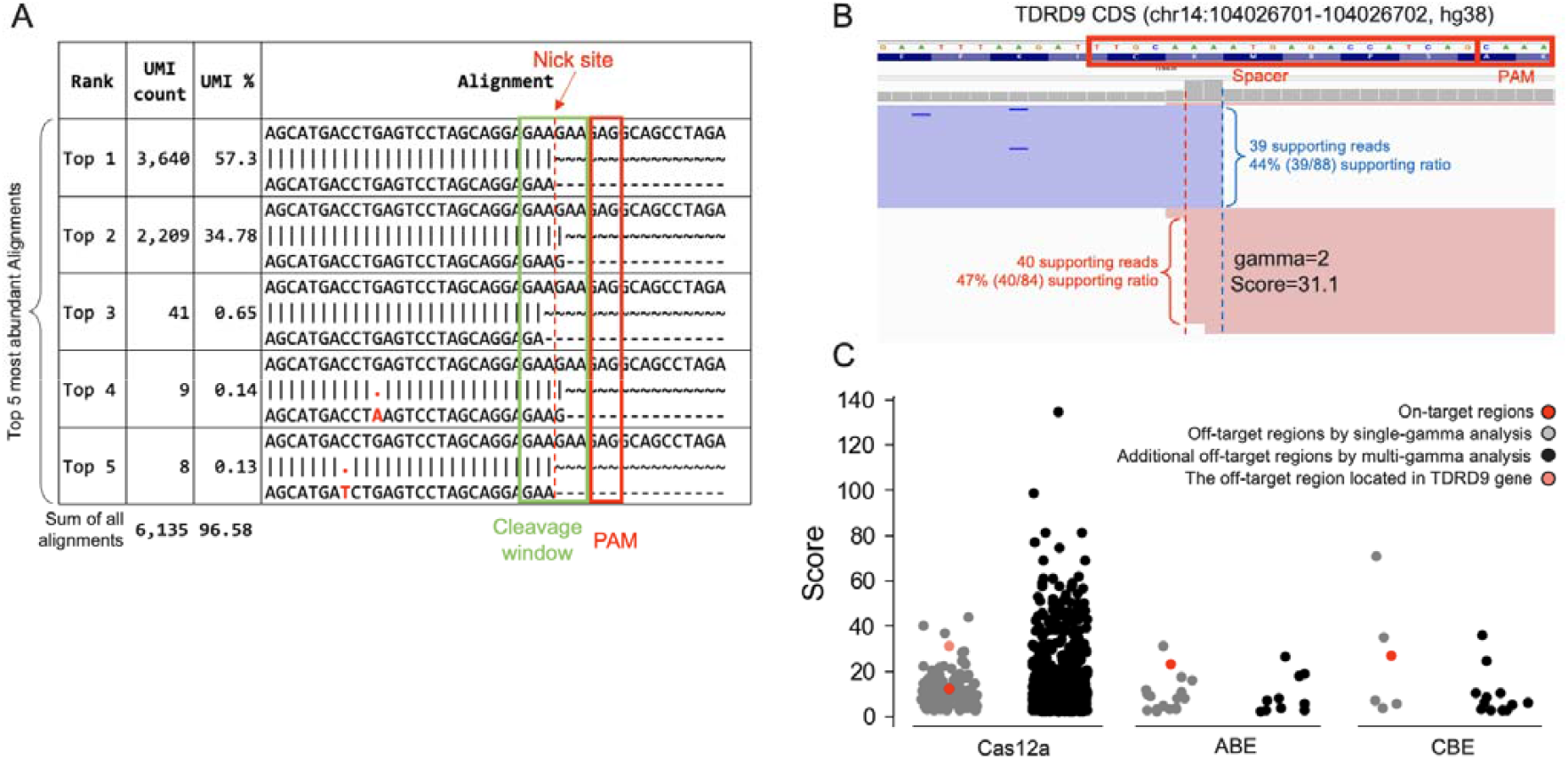
Selected results from SOS and WGS analysis. (A) Alignment of valid and edited UMIs against the proto-side of an Cas9-mediated EMX1 off-target region identified in SOS analysis. The accumulative UMI count and percentage across all valid and edited alignments are indicated beneath the alignment table. Only the sequences of the top 5 alignments with the highest UMI count are shown. For each alignment, reference sequence is shown at the top and observed read at the bottom. (B) Genome browser view of reads mapping in off-target region located in TDRD9 gene resulted from a Cas12a-mediated Digenome-seq data. Reads mapped in forwardly and reversely are shown in salmon and purple colors, respectively. The targeted homology to guide sequence are outlined with red boxes. (C) Score distribution of off-target regions identified by single-gamma analyses and additional off-target regions by multi-gamma analyses. The single-gamma values are 0 (for Cas12a), 11 (for ABE), and 12 (for CBE). The muti-gamma ranges are -10∼10 (for Cas12a) and 1∼19 (for ABE and CBE).

### Identification of cleavage sites from public Digenome-seq data through WGS module

To display the performance and usage of SANTON on the whole genome sequencing-based data, we applied SANTON to identify off-target cleavage sites with scores >2.5 from several representative public datasets treated by different editing systems, including Cas9 [4], Cas12a [10], and base editors such as ABE [11] (Table S2, Additional file 2). SANTON identified 104 cleavage sites from a Cas9-edited dataset targeting HBB gene (ERR1885412) [4], including all four off-targets previously validated using targeted deep-sequencing [4] (Table S3, Additional file 2). For an ABE-edited dataset targeting HEK2 gene (SRR12791392) [11], 27 off-target regions were detected, encompassing all 18 sites reported by the publication study [11] (Table S3, Additional file 2). By fully supporting the analysis of whole genome-based sequencing data from base editors and Cas12a, SANTON fills an important gap in current off-target detection tools, which have been primarily optimized for Cas9. Applying criteria consistent with the original study [10], SANTON identified 242 off-target regions in a Cas12a-edited dataset targeting DNMT1 gene [10], recapturing almost all (40 of the 41) previously reported sites (Table S3, Additional file 2). This high recall rate confirms SANTON’s robust analytical sensitivity, demonstrating its ability to reliably detect true off-target events even in complex whole-genome based sequencing datasets. For the 202 sites additionally identified by SANTON, 74 (36.6%) of them showed cleavage score greater than on-target (score = 12.5). For example, one potential off-target site (score = 31.1) is located within protein-coding region of the TDRD9 gene (Fig. 5B), which encodes a tudor domain-containing protein involved in retrotransposon control in germ cells [12]. Frameshift mutations in exon-5 of this gene has been reported that exists strongly linked to male infertility with azoospermia [13]. These results demonstrate SANTON’s potential to not only reproduce known off-target sites with high accuracy but also discover additional off-target regions missed by prior methods, further establishing its role as a robust and comprehensive computational framework for genome-wide off-target profiling.

### Enhanced features of SANTON for whole genome sequencing-based data analysis

When compared with the latest version of public Digenome tool [6] (Additional file 3), SANTON showed enhanced capabilities for Digenome-seq data analysis while maintaining sensitivity and accuracy comparable to the existing tool in public domain [6] (Fig. S7, Additional file 1). One of key features in SANTON (summarized in Table S4, Additional file 2) is its ability to efficiently process multi-gamma scenarios, particularly for complex cleavage patterns generated by Cas12a and ABE/CBE systems. Unlike the existing Digenome-seq tool [6], which requires separate analytic runs for each gamma value, SANTON explores all gamma values in a single analysis, streamlining the process followed by additional steps including the clustering of nearby sites, redundancy removal, and loci-level summarization (Table S4, Additional file 2). This consolidated approach not only minimizes computational overhead but also ensures more comprehensive detection of potential off-target effects, particularly for base editors where flexible gamma thresholds play a crucial role in identifying subtle editing events. For ABE/CBE systems, SANTON annotates editable adenine or cytosine positions by extracting flanking sequences downstream of the gRNA (Table S4, Additional file 2). SANTON also demonstrates improved computational efficiency, reducing runtime requirements for multi-gamma analyses to approximately ∼2% of traditional workflows (Fig. S8, Additional file 1). This significant reduction in runtime, combined with enhanced sensitivity in off-target identification, highlights SANTON’s efficiency in high-throughput genome-wide analyses, making it a powerful and scalable tool for genome-editing research. This capability enables the identification of additional off-target sites, some with higher scores than the corresponding on-target site (Fig. 5C).

## Conclusions

SANTON is a comprehensive bioinformatics package designed for the identification and quantification of off-target cleavage sites across diverse gene editing platforms, including Cas9, Cas12a, and base editors. By integrating genome-wide sequencing data through its SOS and WGS modules, SANTON enables rigorous analysis of off-target effects. It provides robust support for both Cas9/Cas12a-induced DNA cleavage events and the distinct off-target patterns associated with base editors, which enable the capability that identifying more potential novel off-targeting sites which are missed in the previous study. As of our submission, SANTON is the first reported tool to analyze synthetic oligonucleotide-based sequencing data and, to our knowledge, the first to deploy optimized analytic method for whole genome-based sequencing data generated from Cas12a and base editors. Compared to existing tools, SANTON incorporates advanced features that streamline complex analyses and improve workflow efficiency. Its established framework and modularity allows feasible extension to other off-target nomination assays such as tag-seq [14], site-seq [15] and break-tag [16]. These capabilities establish SANTON as an essential resource for enhancing the precision and reliability of genome editing technologies.

## Availability and requirements

Project name: SANTON

Project home page: https://github.com/VOR-Quantitative-Biology/SANTON

Operating system(s): Platform independent Programming language: Python

Other requirements: none License: Non-commercial license

Any restrictions to use by non-academics: License needed

### List of abbreviations

gRNA: Guide RNA
ABE: Adenine base editor
CBE: Cytosine base editor
NGS: Next generation sequencing
OT: Off-target region
PAM: Protospacer adjacent motif
SOS: Synthetic oligonucleotide-based sequencing
USER: Uracil-specific excision reagent
WGS: Whole genome-base sequencing

## Declarations

### Ethics approval and consent to participate

Not applicable.

#### Consent for publication

Not applicable.

#### Availability of data and materials

Raw NGS sequencing data from EMX1-targeted oligo synthesization based sequencing experiment is available from NCBI GEO (GSE289542). The genome mapping files (BAM files) used for Digenome-seq analysis are available from https://zenodo.org/ with the following IDs: 14736018, 14767914, 14787631, 14788059, 14788344, 14788462, 14791164, and 14791420.

#### Funding

Not applicable.

#### Authors’ contributions

HQ and RW conceived and developed the software. HQ, RW, AM implemented the method and analyzed the data. SW prepared off-target library and generated NGS data. HQ, RW, SW, EA and GG wrote the paper. All authors read and approved the final manuscript.

## Acknowledgements

We thank Michael Pettiglio for valuable discussion of Cas12a biology, Ben Hall for copyright consultant.

