## Supplementary Figures for "SANTON: Sequencing Analysis Toolkits for Off-target Nomination"

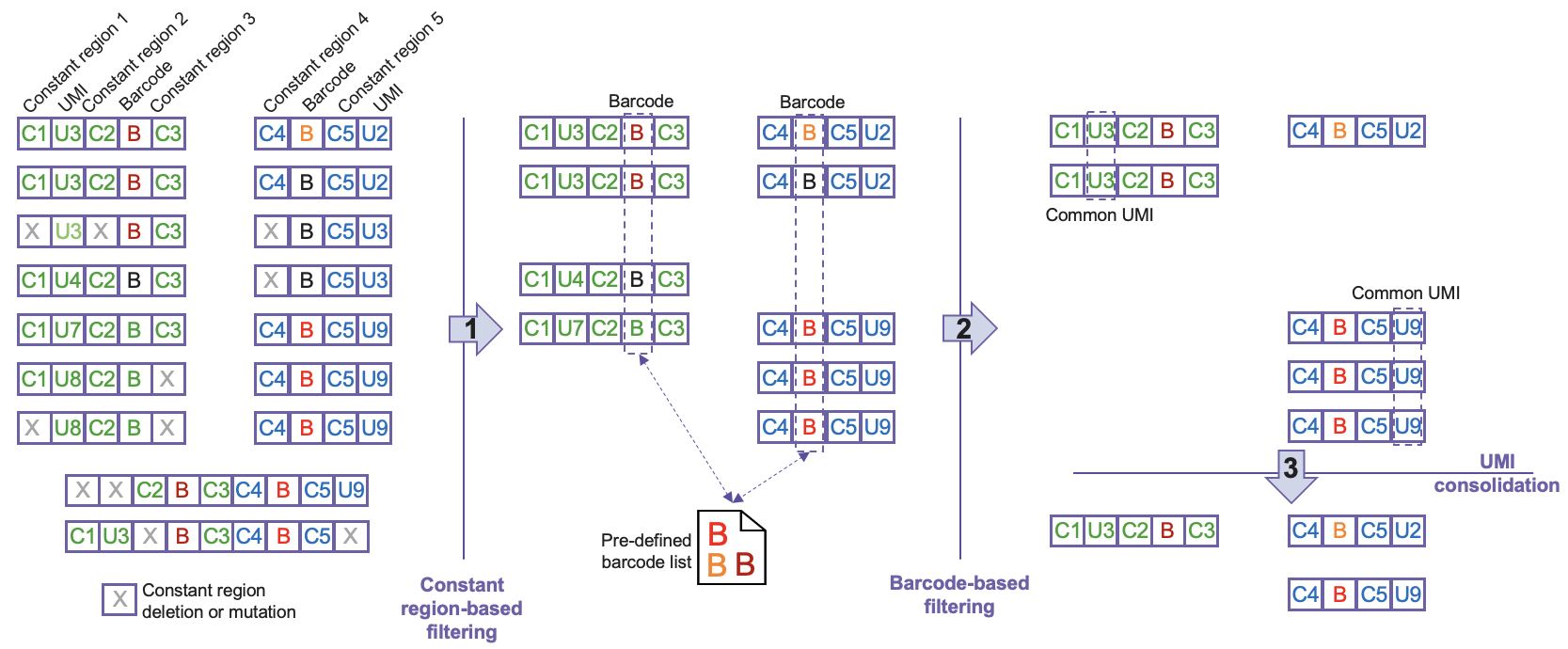

**Fig. S1.** Preprocessing of reads from synthetic oligonucleotide-based sequencing data. 1) Constant region-based data filtering discards pam-side reads without intact constant regions-1, -2, -3. Proto-side reads without intact constant regions-4 and -5 are discarded. Reads containing constant regions from both pam-side and proto-side are discarded. 2) Barcode extracted from pam-side reads and proto-side reads are compared to pre-defined barcode list during library design. Reads with barcode beyond the list are discarded. 3) Reads are clustered based on shared UMIs. For each cluster, the read with the highest quality score is used as a representative sequence.

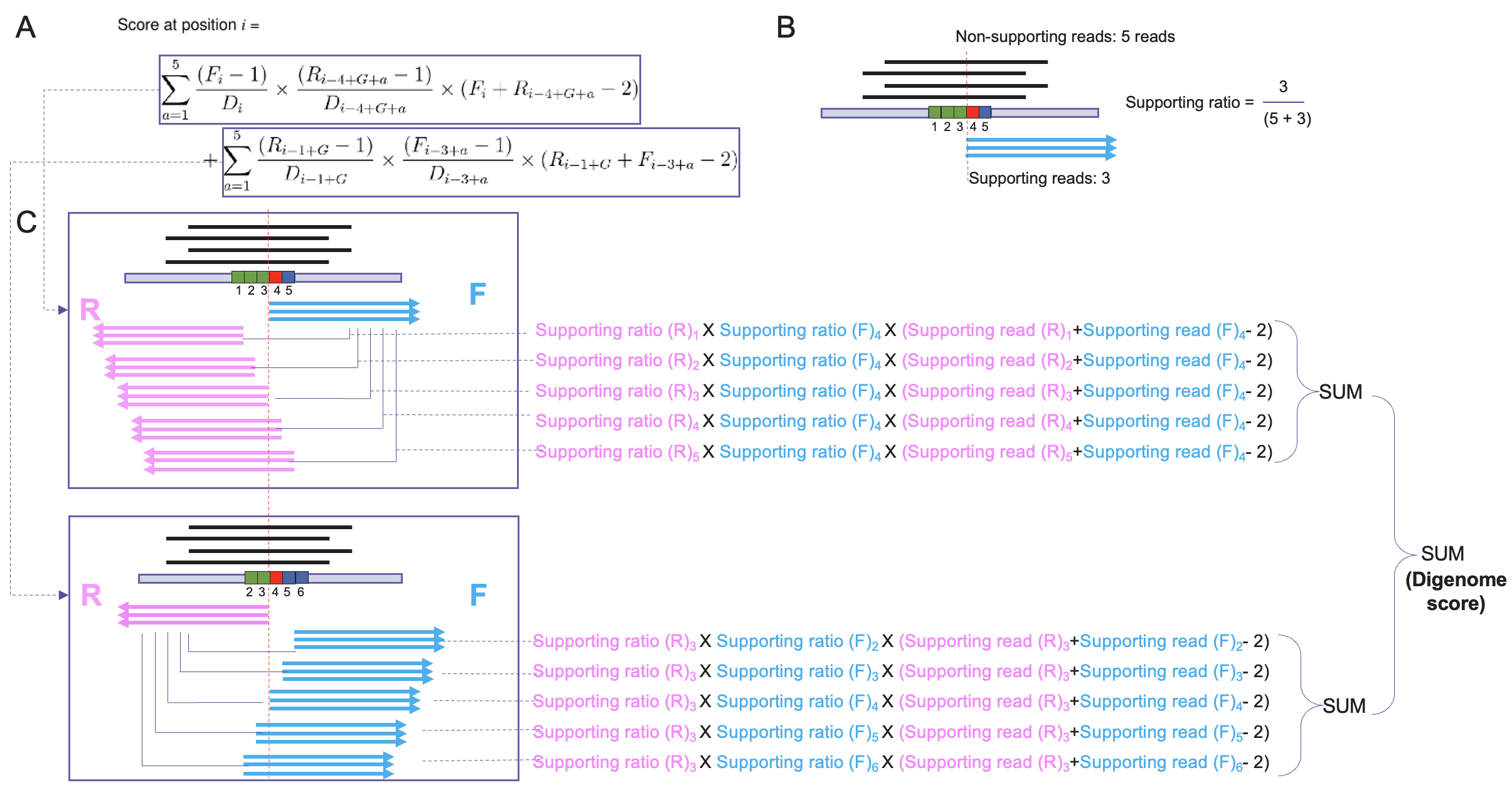

**Fig. S2.** Cleavage score calculation. (A) Cleavage score formulation from previous studies [1, 2]. (B) Definition of supporting ratio. For a given position, the supporting ratio is calculated by dividing the number of supporting reads by the total number of reads covering that position. (C) Schematic illustration of scoring process for a blunt-end cleavage site. Firstly, the weighted supporting ratio is calculated for the 5’-end of read alignment of reverse read cluster as well as its 2bp flanking positions on both sides. For each of these 5 positions, their supporting ratio is multiplied by the supporting ratio of forward read cluster and the sum of supporting read counts from these two read clusters minus 2. Weighted supporting ratios for the 5 positions are summed. Secondly, the weighted supporting ratio for forward read clusters is calculated and summed in a similar way. Lastly, Cleavage score is derived as sum of weighted supporting ratios for forward- and reverse-read clusters.

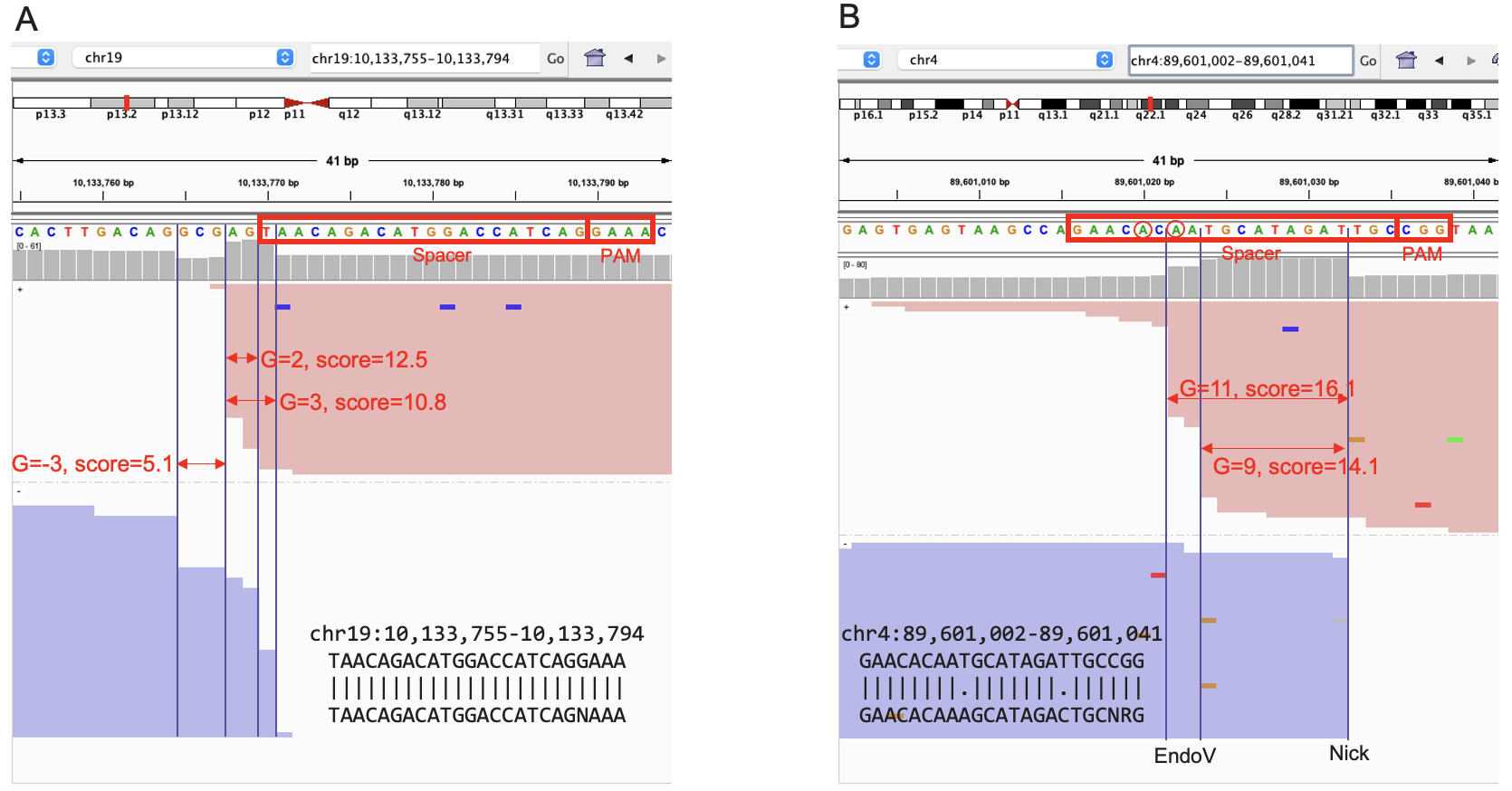

**Fig. S3**. Example of regions containing more than one cleavage site. (A) The on-target region in a Cas12a-treated data with gRNA targeting DNMT1 gene. The top three cleavage sites with the highest score are annotated. The gamma (G) value for each cleavage site is indicated. (B) An off-target region in an ABE-treated data with gRNA targeting HEK2 gene. The two cleavage sites are annotated as earlier mentioned. The editable adenines 5’-upstream to each Endo-V cleavage site are shown in circles.

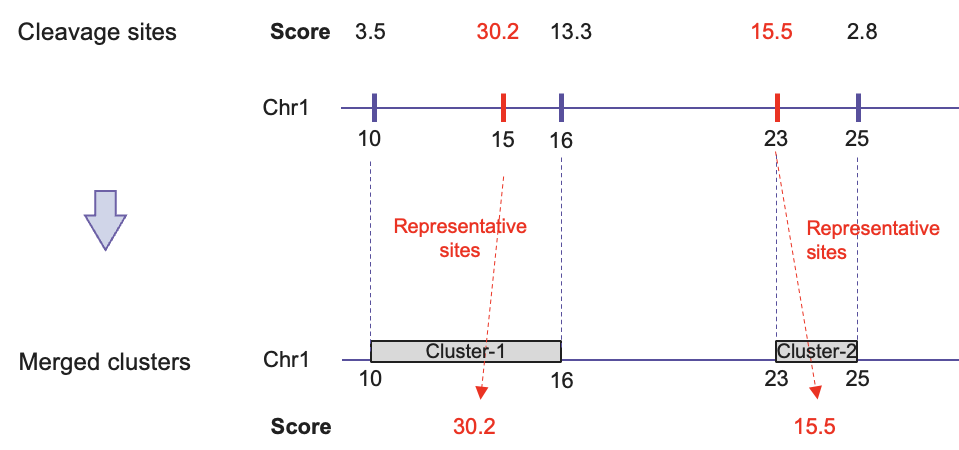

**Fig. S4**. Simplified illustration of cleavage site clustering method. Horizontal lines stand for chromosome 1 reference sequence. Vertical ticks stand for hypothetical cleavage sites with their positions indicated. Red vertical ticks stand for the cleavage sites with the highest score for each cluster. The cleavage sites that are ≤5bp apart are clustered using single-linkage clustering method. The resulting clusters correspond to genomic regions covering all its cleavage sites and are assigned the highest score among its cleavage sites.

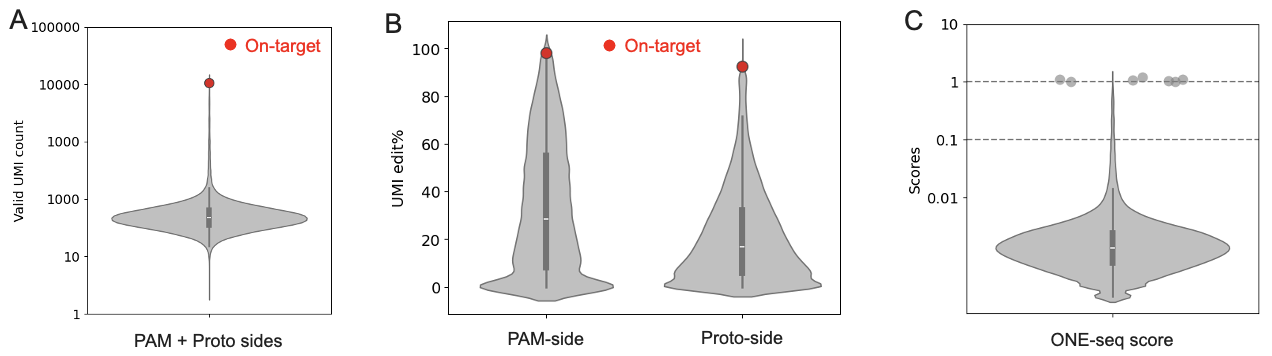

**Fig. S5.** EMX1 off-target data analysis. (A) Distribution of valid UMI counts associated to off-target templates (pam-side and proto-side combined) included in the library. (B) Distribution of edited UMI% associated to examined off-target regions on proto- and PAM-side separately. (C) Distribution of scores across all off-target templates. Off-target regions with score >1 is shown in grey dots.

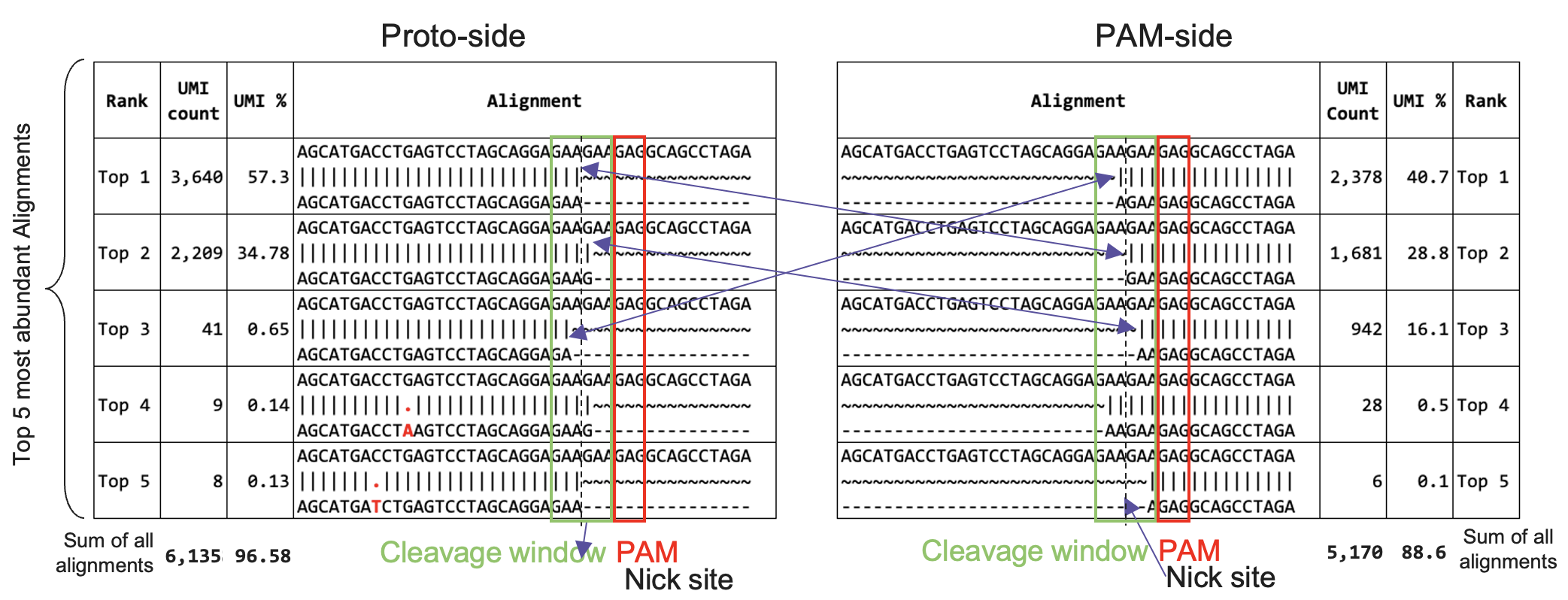

**Fig. S6.** Proto- and pam-side alignments of edited UMIs in a Cas9-mediated EMX1 off-target site identified from synthetic oligonucleotide-based sequencing data. For each side, the accumulative UMI count and percentage across all valid and edited alignments are indicated beneath the alignment table. Only the sequences of the top 5 UMIs with the highest abundance are shown in the figure. Complementary alignments resulted from cleavage at the same nucleotide position are connected by double-arrow lines.

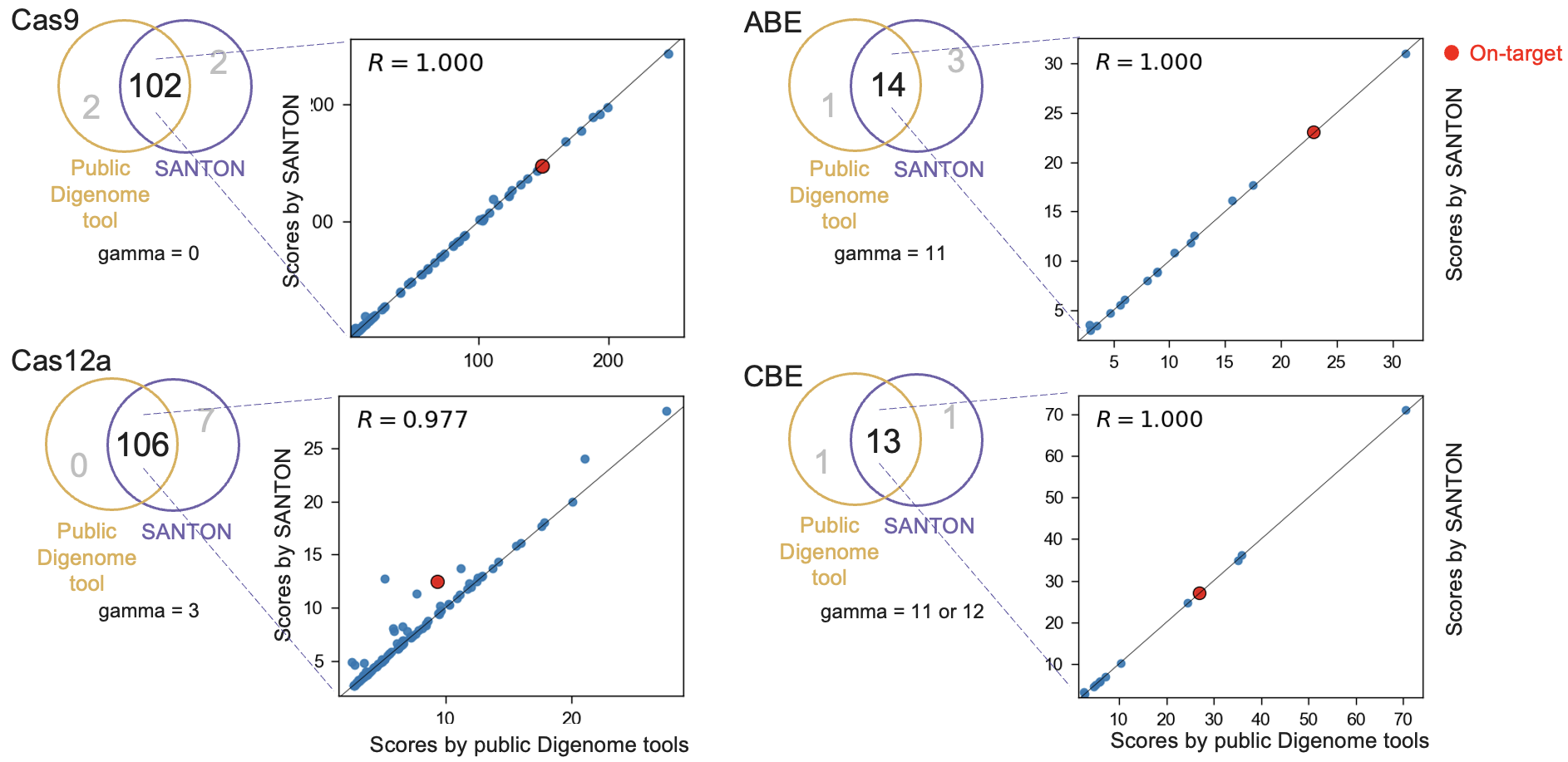

**Fig. S7**. Consistency of cleavage site identification and score between public Digenome-seq tool [2] and SANTON. Four representative datasets were analyzed using the two tools applying the same criterion (see Additional file 3 for details). The shared and unique cleavage sites by the two methods are shown in Venn diagrams. The correlation of scores by the two methods are calculated for share cleavage sites. The gamma values used in analysis of each dataset are indicated (see Additional file 3 for details).

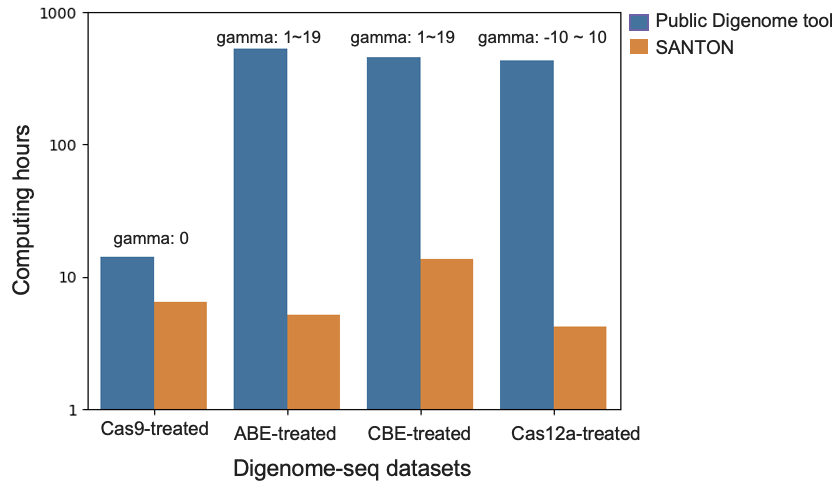

**Fig. S8.** Comparison of computing time used by the public Digenome-seq tool [2] and SANTON for analyses of 4 datasets (Table S2, Additional file 2). Four representative datasets were analyzed using the two tools applying the same criterion (see Additional file 3 for details). The gamma values used in analysis of each dataset are indicated.
