## Supplementary Information for "SANTON: Sequencing Analysis Toolkits for Off-target Nomination"

**Supplementary methods and results**

**EMX1 synthetic oligonucleotide-based sequencing library construction and NGS sequencing**

The protocol used to prepare EMX1 synthetic oligonucleotide-based sequencing library largely followed the published protocol with some modifications [1]. Synthesized EMX1 off-target library was first resuspended in 1X TE buffer to 500 nM. The single-stranded library was amplified and converted to double stranded DNA in a 100 uL PCR reaction with 2 ul of 5 nM library in Q5 high fidelity master mix with primers OS535 and OS536 (Table S5, Additional file 2) at 0.2 uM. The PCR reaction was purified with Ampure beads and the concentration of the eluted library was quantified. For *in vitro* cleavage with SpCas9, the RNP complex was formed at 1:2 molar ratio of Cas9:gRNA and equilibrated at 25 ^o^C for 10 min. The in vitro cleavage reaction consisted of treating 200 ng of library with RNP complex in rNEB3.1 buffer for 2 hours at 37 ^o^C. At the end of incubation, the reaction was treated with 10 ul of Proteinase K (800 U/ml) at 37 ^o^C for 10 min. The reaction was purified with Ampure beads and the treated library was subjected to end extension prior to adaptor ligation. Extended product was prepared in a 50 ul reaction containing 1X Phusion HF buffer, 200 uM dNTPs, 5% DMSO and 1 unit of Phusion DNA polymerase at 72 ^o^C for 10 min. Adaptor was subsequently ligated to the extended product at 25 ^o^C for 5 min after purification. Adaptor ligated product was sized selected for fragments between 100-200 bp using the Bluepippin platform. Size-selected product was purified and amplified by primer OS40, OS101 and OS154 (Table S5, Additional file 2) at 0.5 uM with 200 uM dNTPs, 1 unit of Phusion DNA Polymerase and 1X Phusion HF Buffer. The PCR reaction was purified and diluted 25 fold. Standard barcoding PCR was performed with 1 uM of Illumina barcoding primers and 10 ul of diluted product. The sample was sequenced on MiSeq according to Illumina’s standard protocol.

**Constructing Analysis file for EMX1 synthetic oligonucleotide-based sequencing data analysis**

An analysis file was constructed to include essential metadata for each predicted off-target as well as parameters for data analysis (Table S6, Additional file 2). Briefly, for each off-target (or row), a unique template identifier, on-target status, barcode, template sequence, and coordinate of its genomic region were specified based on the design of sequencing library. For analysis parameters common to all off-targets in a library, the enzyme type is set to ‘Cas9’, expected cut position to 28 (28 bp from 5’-end of template sequence) and extension to 2 (Table S6, Additional file 2).

**EMX1 synthetic oligonucleotide-based sequencing data analysis**

NGS data (fastQ file) derived from cleaved EMX1 off-target library NGS sequencing was trimmed using Trim-galore (<https://www.bioinformatics.babraham.ac.uk/projects/trim_galore/>) with parameters (--paired --quality 20 --phred33 --stringency 4 -e 0.1 --length 80). The trimmed pair-end reads were merged using flash2 [2] with parameters (--min-overlap=15 --max-overlap=300 --max-mismatch-density=0.25 --allow-outies). The resulting fastQ.gz file and analysis file were used as inputs to SOS module in SANTON for bioinformatic analysis.

**Re-analyzing published Digenome-seq data**

The NGS data (fastQ files) from three representative Digenome-seq studies using Cas9 [3], ABE [4], and Cas12a [5] were downloaded from NCBI SRA database under their respective identifiers (Table S2, Additional file 2). The NGS reads were mapped to human reference genome (hg38) using Isaac aligner [6] with settings (--base-quality-cutoff 15 --default-adapters Standard --realign-gaps no --memory-control --variable-read-length yes). The resulting BAM files were analyzed in using WGS module in SANTON. Because no details were provided in these studies [3–5] regarding cleavage site identification, we applied the following parameters to all three datasets: supporting read >5 (forward and reverse), supporting ratio >0.02 (forward and reverse), and score >2.5. The gamma values are 0 for Cas9-treated dataset [3] and 1~19 for ABE-treated dataset [4]. For Cas12a-treated dataset [5], we applied the same additional requirements as the original study did (gamma values: 1~5, and mismatches ≤6bp). The cleavage sites (≤5bp apart) from ABE- and Cas12a-treated datasets were further clustered and merged (Fig. S4, Additional file 1) to remove redundancy. The lists of cleavage sites predicted by SANTON were compared with the equivalents reported in the original studies [3–5]. The cleavage sites appeared in both lists and have genomic coordinates ≤5bp apart are considered shared identifications.

**Accessing accuracy of SANTON WGS module**

We accessed the accuracy of the WGS module in SANTON relative to the latest version of public Digenome tool [3, 7] (<http://www.rgenome.net/digenome-js/standalone>). We used 4 benchmarking Digenome-seq datasets representing cleavages by Cas9, ABE, CBE and Cas12a, respectively (Table S2, Additional file 2). For a fair comparison, we carried out the data analysis at cleavage site-level (without clustering and redundancy removal) and under single-gamma mode. The same criteria were applied while using the two tools: supporting read >5 (forward and reverse), supporting ratio >0.02 (forward and reverse), and score >2.5. The gamma values are 0 for Cas9-treated data, 3 for Cas12a-treated dataset, and 11 for ABE-treated dataset. These gamma values are based on common knowledge for the corresponding editing enzyme and/or learning from alignment pattern observed in on-target region in the corresponding dataset. Because of the scarcity of detected off-target sites in CBE-treated dataset, we used two gamma values (11 and 12) to obtain more datapoints for a meaningful comparison.

The number of off-target sites identified by two tools are highly consistent with most off-target sites being predicted by both tools except minor numbers unique to SANTON (1~7 sites) and to public Digenome tool (0~2 sites) (Fig. S7, Additional file 1). Most unique sites are of low score (Table S7, Additional file 2) and likely reflects artifacts judged by manual inspection. Regarding shared on/off-target sites, their scores from the two tools are nearly identical with exceedingly high correlation (R = 0.97~1, Fig. S7, Additional file 1). These agreements suggest high accuracy of SANTON compared to public Digenome tool [7].

**Assessing SANTON WGS module analysis speed**

We compared SANTON and the latest version of public Digenome tool (<http://www.rgenome.net/digenome-js/standalone>) [7] in terms of analysis speed in an AWS m5.2xlarge EC2 machine with 8 CPUs. For fair comparisons, we applied the same criteria to the two tools: supporting read >5 (forward and reverse), supporting ratio >0.02 (forward and reverse), score >2.5. The gamma values are 0 for Cas-treated dataset, 1~19 for ABE- and CBE-treated dataset and -10~10 for Cas12a-treated dataset. For each dataset, one job is executed for analysis using SANTON under the corresponding analysis mode covering all possible gamma values, and the computing time was recorded. When using the public Digenome tool [7], a job was executed separately for each single gamma value per dataset. The computing time for analyzing a dataset was the sum of computing time used for all gamma values. As shown in Fig. S8 (Additional file 1), the computing time consumed by the two tools for analyzing Cas12a, ABE and CBE each differ dramatically. SANTON used ~2% of time than public Digenome tool [7] primarily due to its support for streamlined multi-gamma analysis.
